# Discovery of shared epigenetic pathways across human phenotypes

**DOI:** 10.1101/2024.04.15.589547

**Authors:** Ilse Krätschmer, Hannah M. Smith, Daniel L. McCartney, Elena Bernabeu, Mahdi Mahmoudi, Archie Campbell, Janie Corley, Sarah E Harris, Simon R. Cox, Riccardo E. Marioni, Matthew R. Robinson

**Affiliations:** Institute of Science and Technology Austria, Klosterneuburg, Austria; Centre for Genomic and Experimental Medicine, Institute of Genetics and Cancer, University of Edinburgh, Edinburgh, UK; Usher Institute, University of Edinburgh, Nine, Edinburgh Bioquarter, 9 Little France Road, Edinburgh, EH16 4UX, UK; Lothian Birth Cohorts, Department of Psychology, University of Edinburgh, Edinburgh, UK

**Keywords:** multi-omics association, EWAS, methylation, prediction, multi-trait

## Abstract

Omics-based association studies typically consider the marginal effects of a feature, such as CpG DNA methylation, on a trait (e.g, independent models for each feature). Although some methods can assess all features together in joint and conditional estimation, this is currently done on a trait-by-trait basis. Here, we introduce MAJA, a method to learn shared and outcome-specific effects for multiple traits in multi-omics data. MAJA determines the unique contribution of individual loci, genes, or molecular pathways, to variation in one or more traits, conditional on all other measured “omics” data genome-wide. Simulations show MAJA accurately finds shared and distinct associations between omics-data and multiple traits and estimates omics-specific (co)variances, allowing for sparsity and correlations within the data. Applying MAJA to 12 outcome traits in Generation Scotland methylation data (n=18,264), we find novel shared epigenetic pathways among cholesterol metabolism, osteoarthritis, blood pressure and asthma. In contrast to marginal testing, we find only 10 CpG probes with significant effects above the genome-wide background. This highlights the need for joint association testing in highly correlated methylation data from whole blood and for studies of increased sample size in order to refine epigenomic associations in observational data.

## Introduction

Epigenetic mechanisms influence gene expression, cell differentiation, tissue development, and disease susceptibility^1,2,3^. Measuring and tracking epigenetic changes through disease progression can provide insight into disease pathogenesis^4^, elucidate environmental and lifestyle factors influencing health, and provide biomarkers for disease diagnosis and risk stratification^5^. To date, most studies have focused on determining the epigenetic basis of traits individually. However, human phenotypes are highly correlated, with shared risk factors and underlying pathways. Estimating the degree to which epigenetic effects are shared across human traits has the potential to reveal shared disease etiology, improve biomarker discovery and maximise outcome prediction.

In genomics, existing methods for the analysis of multiple correlated traits lack flexibility as they: (i) model at most two phenotypes with multiple variance components^6^; (ii) are targeted only for prediction^7^; (iii) fine-map genomic regions independently so that estimates are not conditional on other genome-wide effects^8^; and/or (iv) conduct association testing one variable at a time^9^. Thus, we lack general methods suitable for a range of “omics” data that analyze multiple outcomes jointly, allowing for the inclusion of different data modalities (i.e. methylation, expression, sequence variation, etc.). A predominant focus has been on the estimation of genome-wide correlations^10^, which estimate the degree of similarity in the effects underlying these traits, but do not provide direct insights into specific underlying shared processes. Ideally, we wish to identify individual loci, genes, or molecular pathways that are both shared and unique between traits, and estimate their effects conditional on all other loci, genes, or pathways genome-wide, determining their unique contribution to phenotype. This joint modelling of effects between traits would improve our ability to use multi-modal “omics” data for risk prediction and patient stratification.

Here, we present MAJA, a multivariate multiple linear regression Bayesian joint sparse model. Our Bayesian approach jointly estimates shared effect sizes for multiple traits, potentially across different omics-data, while correcting for correlations within the data and allowing for sparsity. It thus simultaneously finds shared and distinct associations between omics-data and multiple traits, and estimates group-specific (co)variances. It is scalable, flexible, and suitable for all existing high-dimensional genomics data. We demonstrate our approach using the Generation Scotland data^11^, a cohort of 18,264 individuals with blood-based methylation measures, where we find both trait-specific and shared probe effects and improve out-of-sample prediction as compared to single-trait models.

## Multivariate joint regression model: MAJA

We developed a multivariate Bayesian multiple regression model (MAJA MultivAriate Joint bAyesian model) that jointly estimates omics effects and their corrections on multiple traits and performs variable selection, all while taking into account correlations within the data. Our model is suitable for the case where a number of *q* phenotypes for *n* individuals is measured within the matrix ***Y***. The phenotype matrix is modelled to be linearly related to the matrix containing *p* genomics measures ***X*** (e.g. SNPs, epigenetic probes, gene expression) as,

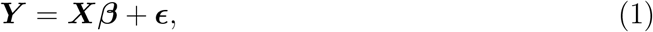

where the matrix ***β*** denotes the effect sizes for *q* traits and ***ϵ*** represents the residual error matrix. Each column of the ***Y*** and ***X*** matrices is standardized. The parameters of Equation 1 are estimated using a Gibbs sampler, an iterative Markov Chain Monte Carlo method. All details on MAJA can be found in the Materials and Methods section.

The effects of each genomic location *j* on the multiple traits, ***β***_***j***_, are assumed to have a multivariate spike-and-slab prior distribution to accommodate zero effects sizes

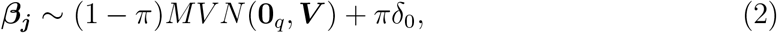

where *π* is the probe exclusion probability common to all traits, *MV N* (**0**_*q*_, ***V***) is a multivariate normal distribution with mean 0 and (co)variance ***V*** and *δ*_0_ the Dirac delta distribution. Through sampling the effects of each probe conditional on the other probes, correlations between the probes are automatically taken into account in our model.

Moreover, MAJA is able to handle multiple ***X*** matrices. For example: (i) (epi)genetic information split into groups where the (epi)genetic covariances are estimated within each group; and/or (ii) multi-modal data, where different data sets are combined, like CpG sites and single nucleotide polymorphisms (SNPs). Effect sizes are determined jointly, thus the effects each column of ***X*** are estimated conditional on all others, taking into account correlations across groups and omics layers.

We demonstrate that MAJA accurately infers (co)variances in one or multiple groups using simulations as described in the Materials and Methods and shown in Figures S1 and S2. We also show that the estimated effects can be used to predict into a test data set, to achieve out-of-sample prediction accuracy that conforms to theoretical expectations and improves over single-trait models, as can be seen in Figures S3 and S4. Finally, we demonstrate the ability of MAJA to localise effects accurately to the single-variable level, conditional on all other variables, by calculating the true positive (TPR) and false discovery (FDR) rate across all simulation scenarios, displayed in Figure S5.

### Multi-trait epigenetics in Generation Scotland

We apply MAJA to 18,264 individuals in Generation Scotland for whom DNA methylation measures from whole blood were available at 831,349 CpG sites for twelve outcome traits, split into six cognitive, two metabolic and four disease traits. For disease outcomes that were commonly self-reported at the time of blood sampling, the phenotypic variance attributable to the methylation probes ranged between 24% for both depression and asthma, to 69% for hypertension. For clinically measured variables, we find that the phenotypic variance attributable to the methylation probes was 81% for body mass index (BMI) and 73% for ratio of high density lipoprotein over total cholesterol. In addition, we extend our analysis to a series of cognitive evaluations and educational attainment metrics, finding that between 32% and 73% of the phenotypic variation can be attributed to the CpG probes. The estimated variances, covariances between the traits and correlations are shown in Figures 1 and 2. The values along with the 95% credible intervals are listed in Supplementary Tables S1-S4. Heatmaps of the correlations can be found in Supplementary Figure S6.

**Figure 1:**
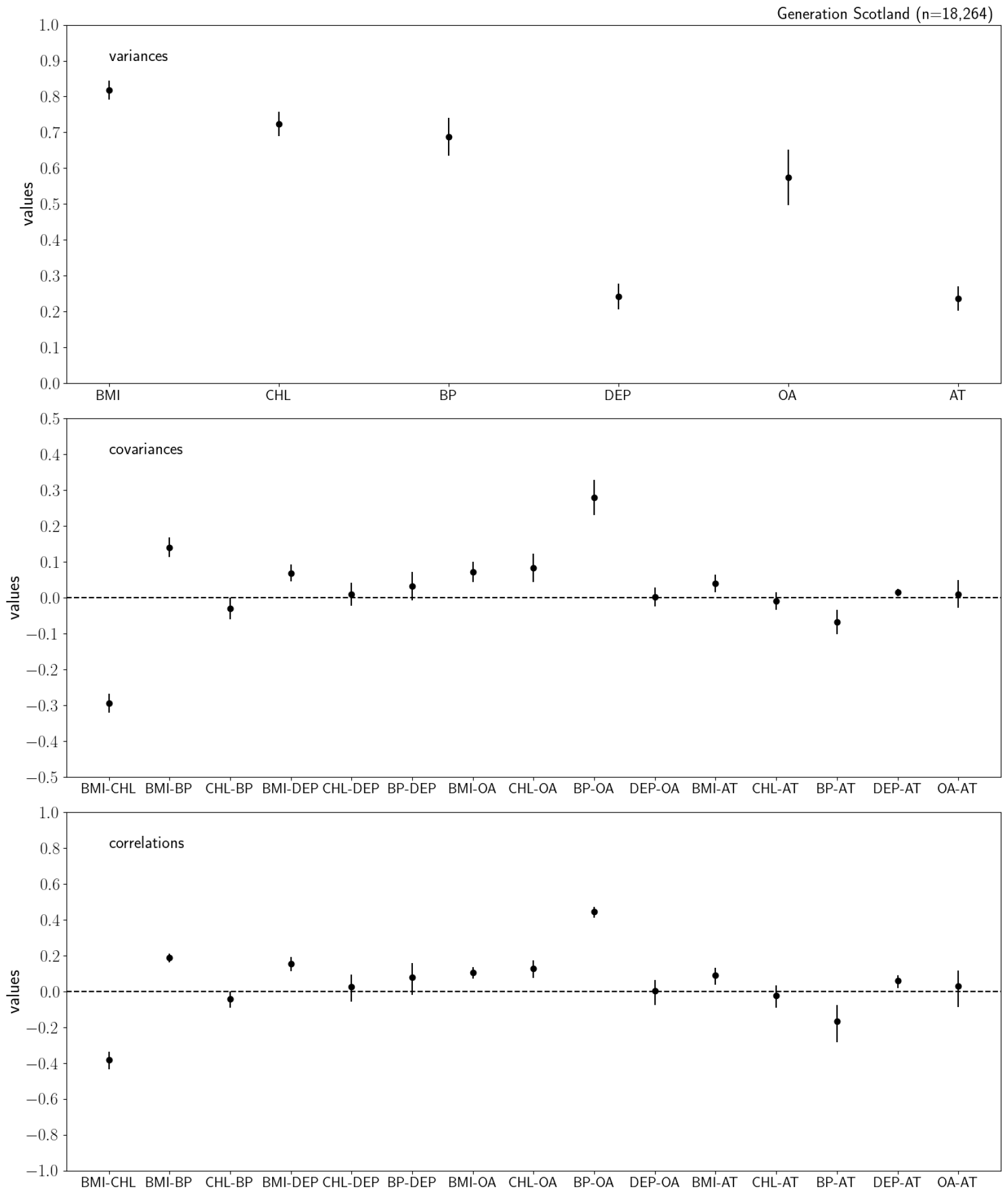
Estimated epigenetic variances (top), covariances (middle) and correlations (bottom) for body mass index (BMI), ratio of high density lipoprotein over total cholesterol (CHL), high blood pressure (BP), depression (DEP), osteoarthritis (OA) and asthma (AT) in the Generation Scotland methylation data using 18,624 individuals and 831,349 probes. The error bars in the upper plot represent the 95% credible interval. The correlations are calculated as covariances scaled by the corresponding variances. The uncertainties are calculated using the posterior means *±* 95% credible interval.

**Figure 2:**
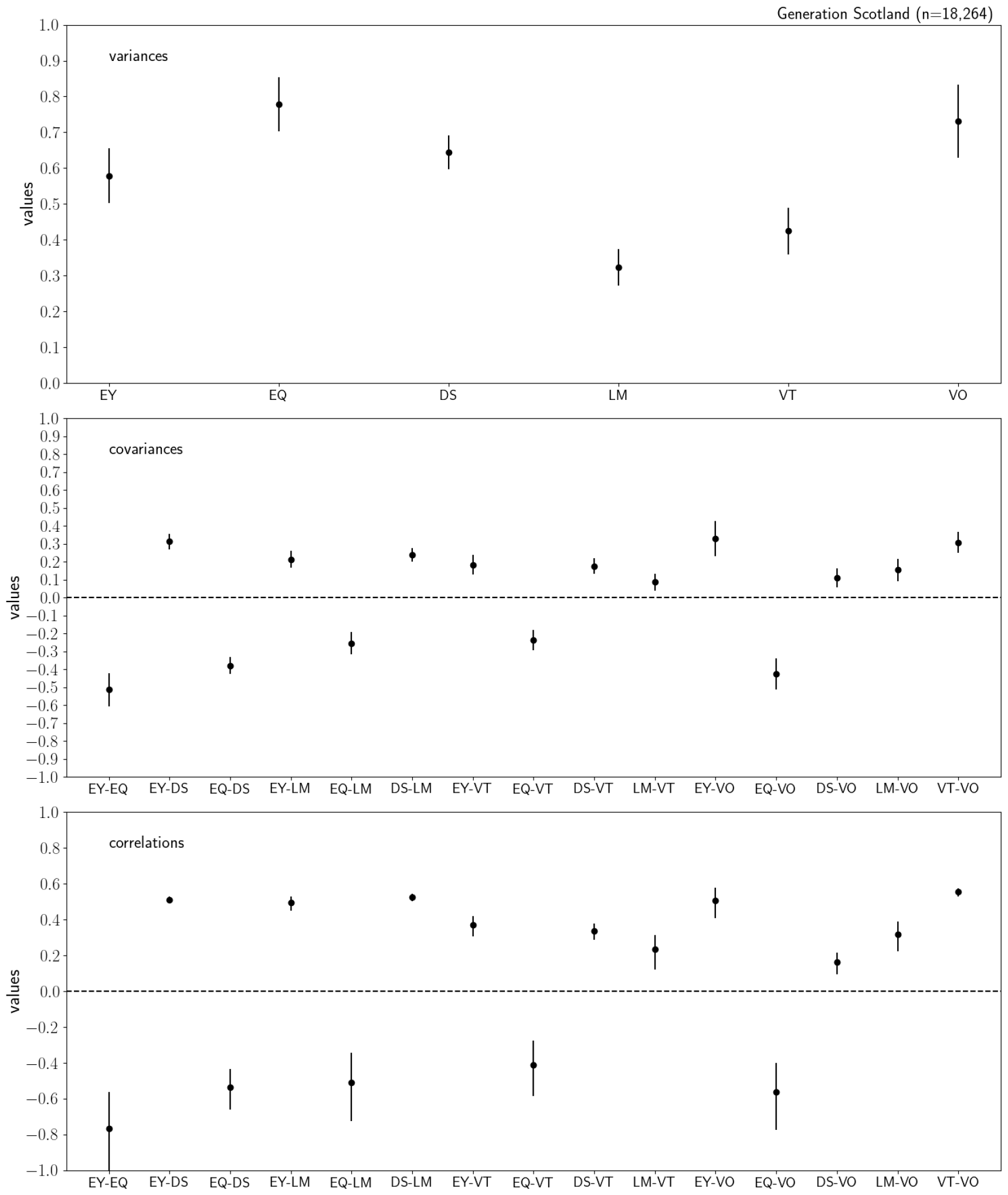
Estimated epigenetic variances (top), covariances (middle) and correlations (bottom) for years in education (EY), highest qualification in education (EQ), digit symbol (DS), logical memory (LM), verbal fluency (VT) and vocabulary (VO) tests in the Generation Scotland methylation data using 18,624 individuals and 831,349 probes. The error bars in the upper plot represent the 95% credible interval. The correlations are calculated as covariances scaled by the corresponding variances. The uncertainties are calculated using the posterior means *±* 95% credible interval.

We find a strong negative correlation of epigenetic effects between BMI and ratio of high density lipoprotein over total cholesterol and that CpG probe effects were positively correlated for BMI and all other traits. CpG effects for both ratio of high density lipoprotein over total cholesterol and self-reported hypertension are positively correlated with those for self-reported osteoarthritis. Interestingly, CpG effects for self-reported asthma are negatively correlated with those for hypertension, implying asthma-associated epigenetic probes have an inverse association for hypertension.

We find weak correlations of epigenetic effects among digit symbol and vocabulary cognitive tests, but generally strong correlations among all other tests. Cognitive tests share underlying methylation probe effects with both years of education and educational attainment. Note here that the highest educational attainment is scored as a “1” (see Materials and Methods) and thus the negative correlation reflects methylation effects acting in the same direction for longer years in education and higher educational attainment.

There are no strong residual correlations, as can be seen from Figures S7 and S8, which implies that methylation probe variation captures the vast majority of the signal of trait correlations. Residual covariances that are non-zero are often in contrast to the methylation covariance, implying relationships among risk factors not captured by methylation patterns in whole blood differ to those reflected in the covariance of methylation probes effects.

We find nine unique probes whose effects, conditional on those of all other probes, have an posterior inclusion probability (PIP) above 95%. MAJA is designed such that a probe or locus affects all the traits or none of them, but the estimated effect size is allowed to differ for each trait, e.g. it can be zero for some traits. Thus, to detect significant associations we use the posterior distribution of effect sizes to determine which of the estimated effects do not include 0 within either one or two standard deviations. Using both the inclusion probability and the strength of the effect size provides a robust test statistic for fine-mapping CpG effects, as shown in Figure S5. A list of the probes, their related genes and their associated traits is given in Table 1.

**Table 1:**
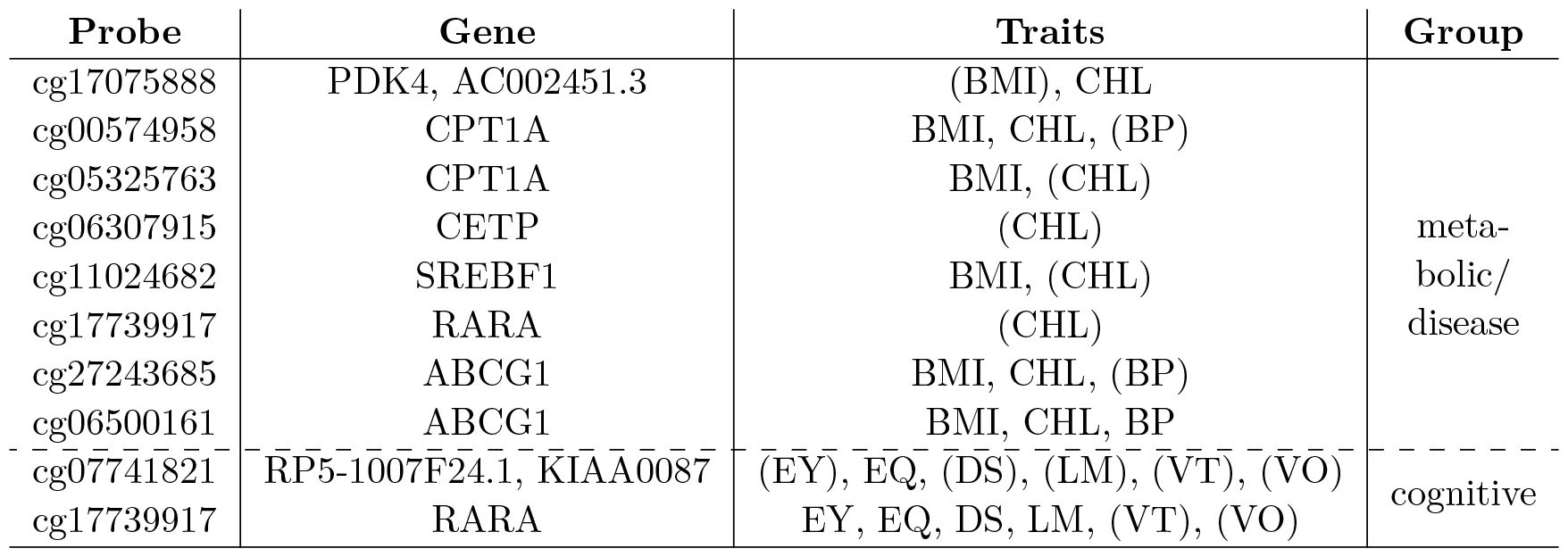
Associated probes with inclusion probability *≥* 95%. Traits listed are those where the effect size of the probe does not include 0 within their standard deviations (SD). Traits tested with 1SD are in brackets, 2SD without brackets.

| Probe | Gene | Traits | Group |
| --- | --- | --- | --- |
| cg17075888 | PDK4, AC002451.3 | (BMI), CHL | meta-<br>bolic/<br>disease |
| cg00574958 | CPT1A | BMI, CHL, (BP) |  |
| cg05325763 | CPT1A | BMI, (CHL) |  |
| cg06307915 | CETP | (CHL) |  |
| cg11024682 | SREBF1 | BMI, (CHL) |  |
| cg17739917 | RARA | (CHL) |  |
| cg27243685 | ABCG1 | BMI, CHL, (BP) |  |
| cg06500161 | ABCG1 | BMI, CHL, BP |  |
| cg07741821 | RP5-1007F24.1, KIAA0087 | (EY), EQ, (DS), (LM), (VT), (VO) | cognitive |
| cg17739917 | RARA | EY, EQ, DS, LM, (VT), (VO) |  |

None of the probes with *PIP≥* 0.95 within the metabolic and disease traits group are associated with depression, osteoarthritis or asthma. Of the seven we discover, almost all are shared between BMI, ratio of high density lipoprotein over total cholesterol, and hypertension. These probes are located near *ABCG1* which controls lipoprotein lipase (LPL) activity and promotes lipid accumulation in human macrophages in the presence of triglyceride-rich lipoproteins; *CPT1A* which is the gatekeeper enzyme for mitochondrial fatty acid oxidation; and *PDK4*, a regulator of pyruvate dehydrogenase (PDH), which influences acetyl-CoA from beta-oxidation into the citric acid (TCA) cycle, thereby leading to enhanced fatty acid (FA) oxidation and slowing of glycolysis or glycolytic intermediates to alternative metabolic pathways. We then additionally find three genes linked to ratio of high density lipoprotein over total cholesterol: *CETP* which is a hydrophobic plasma glycoprotein that mediates the transfer and exchange of cholesteryl ester and triglyceride between plasma lipoproteins, playing an important role in high density lipoprotein metabolism; *SREBF1* which regulates the uptake and synthesis of cholesterol; and *RARA* a key regulator of lipid/glucose metabolism. All of these associations have been reported in the epigenome wide association study (EWAS) catalogue for these, or related, traits; however, here we are able to explicitly determine for which traits their effects are shared and for which they act in a trait-dependent manner and to show that the association holds conditional on all other methylation loci.

Interestingly, the methylation effects of probe cg17739917 near *RARA* which is associated with ratio of HDL over total cholesterol is also linked to all cognitive tests. Also cg07741821 near genes *RP5-1007F24*.*1, KIAA0087* is associated with variation in all cognitive traits. These two associations have not been reported before and taken together, our results show that key pathways are identified by our model whose effects are determined conditional on the data structure and all other probe effects.

Finally, we wished to demonstrate that our approach facilitates improved out-of-sample prediction as compared to single-trait approaches. Taking the CpG effects estimated in Generation Scotland, we predict traits that were measured in the Lothian Birth Cohort (LBC) 1936^12^. We find that multi-trait predictors generally outperform the comparable single-trait predictors calculated using the BayesR model^5^, as shown in Table 2. Of particular note is the predictor of general cognitive function, which explained up to 8.6% of the variance. This is more than double the performance of a previous predictor, derived from a subset of the Generation Scotland dataset^13^.

**Table 2:** Out-of-sample prediction accuracy of episcores (incremental test *R*^2^) created from MAJA as compared to estimates made using the single-trait BayesR model. Episcores were created for a given trait (“Episcore”), using CpG probe estimates from either single trait (“Univariate”), or multitrait (“Multivariate”) models. The episcores were then used to predict a series of outcome traits (“Traits”) in the Lothian Birth Cohort (LBC) 1936 study (n=861). The *R*^2^ values give the incremental test *R*^2^ of including the episcore in a linear model to predict each outcome, adjusting for age and sex. WTAR refers to the Wechsler Test of Adult Reading; NART to the National Adult Reading Test; BMI to body mass index; CHL to ratio of high density lipoprotein over total cholesterol in whole blood. All scores refers to the variance explained by including all predictors together within the model.

| Trait | Episcore | $R^2$ | |
| --- | --- | --- | --- |
|  |  | BayesR<br>Univariate | MAJA<br>Multivariate |
| BMI | BMI | 14.94 | 16.86 |
| BMI (log) | BMI | 14.84 | 16.8 |
| CHL | CHL | 7.02 | 5.06 |
| Digit symbol | Digit symbol | 3.17 | 3.35 |
| Logical memory | Logical memory | 0.47 | 1.06 |
| Verbal fluency | Verbal fluency | 0.88 | 0.74 |
| Years in education | Years in education | 4.10 | 5.32 |
| WTAR | Vocabulary | 3.72 | 4.03 |
| NART | Vocabulary | 4.62 | 4.68 |
| General cognitive function | Logical memory | 3.2 | 5.9 |
| General cognitive function | Vocabulary | 5.2 | 5.5 |
| General cognitive function | Verbal total | 2.2 | 4.8 |
| General cognitive function | Digit symbol | 5.2 | 6.7 |
| General cognitive function | Education years | 6.3 | 7.6 |
| General cognitive function | All scores |  | 8.6 |

## Discussion

We presented MAJA, a Bayesian method that jointly estimates the effect sizes of (epi)genomic variants, as well as correlations of the effects for multiple traits, while correcting for correlations among variables and allowing for sparsity. We extend previous studies both in terms of methodology and in the phenotypes studied. The variance estimates obtained for BMI agree with previous estimates^5^, as does our finding of a strong negative correlation of epigenetic effects between BMI and ratio of high density lipoprotein over total cholesterol^4^. We highlight novel CpG covariances among hypertension and osteoarthritis, ratio of high density lipoprotein over total cholesterol and osteoarthritis, BMI and hypertension, and BMI and asthma, and our results imply that methylation patterns of cholesterol metabolism related genes in whole blood are associated with osteoarthritis pathogenesis and that there is potentially a complex, yet to be fully explored, relationship between hypertension and asthma.

In this work, we focused on developing a statistical model and associated software, and to give a demonstration of how multi-trait Bayesian models can improve the discovery of shared loci and external trait predictions. There are several limitations to our study, mainly the sample size of Generation Scotland, which whilst representing one of the largest single cohorts with methylation data available, still has very limited power to detect associations at *≥* 95% confidence and to produce high out-of-sample accuracy, relative to the estimated total variance attributable to all CpGs on the array. Our model will likely return fewer associations than standard one-probe-at-a-time significance testing. However, single probe analyses do not control for correlations across probes and effects are not estimated conditional on all other probes effects. Thus, single-probe testing likely gives estimates that are inflated by correlations and by general data structure and confounding. In contrast, effect sizes and significance are determined jointly within MAJA which we expect (and show in simulation) to provide an accurate determination and localisation of specific probe effects.

Note that, within our model, a probe will be included for all traits when it has an effect on at least one of the traits. However, the estimates for each trait are freely sampled from a multivariate normal with zero mean and thus there is no reason to expect a directional bias in the effect size estimates for traits for which the probe is not associated. However, estimation error may increase if many small effects are sampled for traits with strong covariance (BMI-ratio of high density lipoprotein over total cholesterol, for example) and this is likely the reason for the loss of out-of-sample accuracy, which we see for the multitrait predictor of ratio of high density lipoprotein over total cholesterol in the LBC1936. This is a modelling choice to facilitate improved association testing and can be overcome by simply setting the effects of these probes to zero if the posterior estimate includes zero within the standard deviation.

Additionally, we highlight that CpG measures in whole blood do not necessarily represent the correct tissue for understanding mechanistic pathways among outcomes. A full characterisation of methylation across multiple tissues is needed to fully capture these relationships. With increasing cross-tissue data, we expect that the ability of MAJA to fit multiple groups could be useful, where multiple cross-tissue methylation measures could be fit within the model to determine patterns of shared effects across tissues. Further limitations are that while MAJA is able to handle larger biobank scale data sets through the use of message-passing interface (MPI) coding, at present it remains computationally expensive and is set up in such a way that all data needs to fit into RAM. Having demonstrated the effectiveness of this framework, our future work will now focus on alternative algorithms for joint inference from this model within a genomics setting.

In summary, our approach provides a method to learn shared and trait-specific epigenetic pathways and to improve prediction of outcomes from omics data. Our approach can be used in future to understand the multi-stage transition from a “pre-disease” to “disease” state, with the overall goal of improving primary prevention, patient stratification, and subsequent clinical management.

## Materials and methods

### Statistical model

Consider n individuals with q observed phenotypes and p recorded (epi)genetic markers. The relationship between the phenotype matrix, ***Y***, of dimensions (n x q) and the design matrix, ***X***, of dimensions (n x p) is modelled as

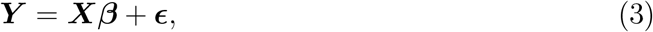

where the (p x q) matrix ***β*** represents the effect sizes while ***ϵ*** is the residual error matrix with dimensions (n x q). The design matrix can be split into various groups according to biological annotations. The design and phenotype matrix are standardized for each column.

We assume that ***Y*** is distributed like a matrix normal with mean ***Xβ***, among-row variance 𝕀_*n*_ (where 𝕀_*n*_ is the unitary matrix with dimension (n x n)) and among-column variance **∑** with dimension (q x q):

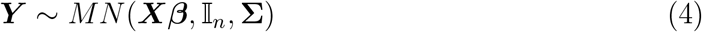

The matrix normal distribution is related to the multivariate normal as

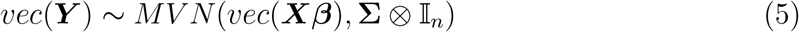

where *vec*(***Y***) represents the vectorization of ***Y*** and ⊗ the Kronecker product. The prior distribution for the residual error matrix ***ϵ*** is also assumed to be a matrix normal *MN* (**0**_(*n*x*q*)_, 𝕀_*n*_, **∑**).

The effects of each marker j on the multiple traits, ***β***_***j***_, is modelled as a multivariate normal with mean **0** and variance ***V***_***g***_ of dimensions (q x q):

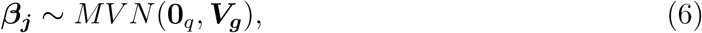

where g refers to the group the marker is attributed to. The group variance ***V***_***g***_ is specific to each group. To be able to model sparsity in the effects, the Dirac delta *δ*_0_ is included in the prior distribution of ***β***_***j***_ with the prior group-specific exclusion probability *π*_*g*_:

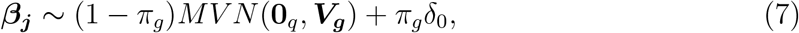

where *π*_*g*_ is modelled by the Dirichlet distribution.

Covariances ***V***_***g***_ as well as **∑** (jointly denoted as ***cov***) are modelled as outlined in Section 2 of Ref.^14^, using a modified Cholesky decomposition:

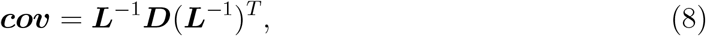

Where

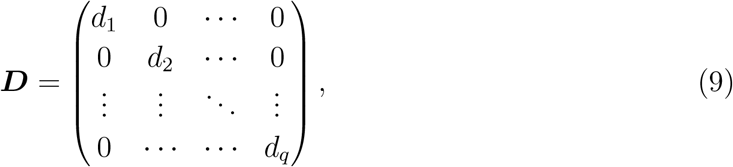

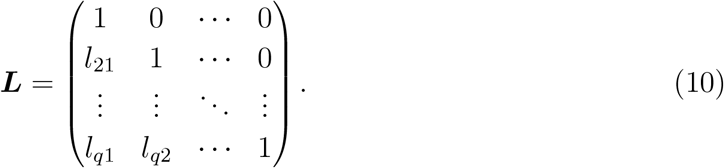

This parameterisation of the variance matrices is advantageous compared to the inverse Wishart distribution (which is the conjugate of the multivariate normal distribution) as the elements in L are unrestricted. They are modelled with a multivariate normal distribution with prior mean 0 and variance *s*_0_ = 0.0001. The prior distribution of the diagonal elements of ***D***, which have to be positive, are set to an inverse Gamma distribution *G*^*−*1^(*a/*2, *ab/*2), where *a* and *b* are the prior shape and scale parameter of the inverse Gamma distribution (here *a* = 2 and *b* = 0.1).

### Gibbs sampler

To estimate the unknown parameters in Equation 3, a Gibbs sampler which is a Markov Chain Monte Carlo (MCMC) method, is set up. The Gibbs sampler runs the following steps for a chosen number of iterations:

1. Sample intercept from a normal distribution.
2. Randomly pick a marker *j* and sample ***β***_***j***_ from its conditional posterior distribution

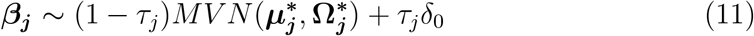

with posterior covariance

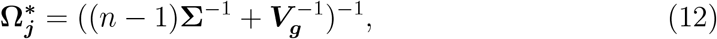

posterior mean

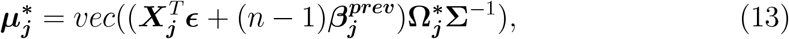

With 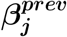 referring to the effects of marker *j* in the previous iteration, and posterior exclusion probability

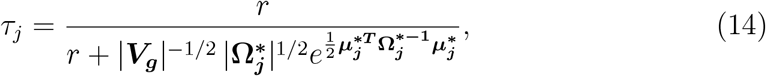

Where 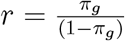.
3. Repeat step (2) until all markers are sampled.
4. Sample exclusion probabilities *π*_g_ for each group from *Dirichlet*(*p*_*g*_ *− Z*_*g*_, *Z*_*g*_), where *p*_*g*_ is the total number of markers and *Z*_*g*_ is the number of non-zero markers in each group.
5. Calculate 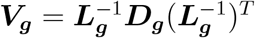 for each group by:
  a. Sampling the diagonal elements of ***D***_***g***_ from

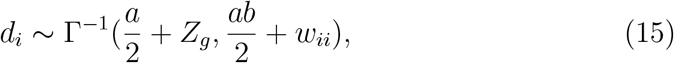

where *w*_*ii*_ is element (i,i) of 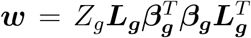. When sampling with more than one group, *Z*_*g*_ and ***w*** are group-specific.
  b. Sampling the elements of the lower triangular matrix ***L***_***g***_ from a multivariate normal distribution with mean

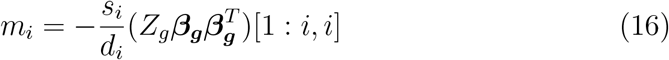

and variance

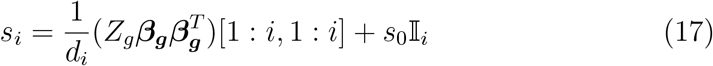

for each row i of ***L***_***g***_ where [1 : *i*, 1 : *i*] denotes the submatrix of between rows 1 to i and columns 1 to i and *s*_0_ is the initial variance of the multivariate normal.
6. Calculate the covariance **∑** in the same way as ***V***_***g***_.

The effects of the markers are sampled conditional on all the other markers, thus automatically taking into account correlations betweeen markers or linkage disequilibrium (LD). The means and variances of ***β, V***_***g***_ and **∑** averaged across iterations (excluding the results from the burn-in period) are stored.

The Gibbs sampler is run for in total 5000 iterations, whereof 1000 are discarded as burn-in. The burn-in period of 1000 is chosen to be well away from the point where the sampler first reaches convergence to make sure that the values for posterior means are only taken when the estimates are stable. Multiple chains are run after the burn-in period for the estimation of the posterior means.

The code requires as input a phenotype matrix without any missing values and a standardized design matrix which has to fit into RAM. For further details on the implementation of the sampler using a Bulk synchronous parallel Gibbs sampling scheme with message passing interface^15^, see links in Code availability.

### Generation Scotland methylation data

Generation Scotland is a large population-based, family-structured cohort of over 24,000 individuals aged 18–99 years^11^. The study baseline took place between 2006 and 2011 and included detailed cognitive, physical, and health questionnaires, along with sample donation for genetic and biomarker data.

The Generation Scotland methylation (GSM) data includes cytosine-phosphate-guanine dinucleotides (CpG sites) for 18,413 individuals. DNA methylation data were processed and quality-controlled in four batches, following broadly similar procedures. Probe and sample quality was assessed using the meffil package in R^16^. Probes were excluded based on low detection P-value (*≥* 0.5% of samples with detection P *≥* 0.05 [batch 1]; *≥* 1% of samples with detection P *≥* 0.01 [batches 2-4]) and low bead count (*<* 3 in *>* 5% of samples). Samples were removed based on 1) a high proportion of probes with high detection-P-values (*≥* 1% of CpGs with detection P-value *≥* 0.05 [batch 1]; *≥* 0.5% of CpGs with detection p-value ≥ 0.01 [batches 2-4]), 2) where recorded sex did not match predicted sex based on information from sex chromosomes, and 3) outlier values based on log median intensites of methylated vs unmethylated signals. The four quality-controlled batches were normalised as a single dataset using the dasen method in wateRmelon^17^. The final set of 831,349 methylation probes were then adjusted for age, sex, smoking and batch and standardized to mean zero and variance one.

We jointly analyze the following phenotypes: body-mass-index (BMI *kg/m*^2^); ratio of high density lipoprotein over total cholesterol (CHL, both measured in mmol/L); selfreported high blood pressure (BP, 2472 cases); self-reported depression (DEP, 1807 cases); self-reported osteoarthitis (OA, 1355); self-reported asthma (AT, 2097 cases); logical memory (verbal declarative memory), calculated from the Wechsler Logical Memory test by taking the sum of immediate and delayed recall of one oral story^11^; digit symbol, ascertained from the Wechsler Digit Symbol Substitution test in which participants recoded digits to symbols over a 120 second period^11^; verbal fluency phenotype, measuring executive functioning, was derived from the phonemic verbal fluency test, using the letters C, F and L, each for 1 min^11^; vocabulary, measured using the Mill Hill Vocabulary Scale, junior and senior synonyms combined^11^; year spent in education; and finally highest educational qualification achieved. Years spent in education was self-reported as the total years attended school/study full-time, with coding 0: 0, 1: 1-4,2: 5-9, 3: 10-11, 4: 12-13, 5: 14-15, 6: 16-17, 7: 18-19, 8: 20-21, 9: 22-23, 10: more than 24 years. For highest educational qualification participants were asked what the highest educational qualification they have obtained, with data then coded as: 1 College or University degree, 2 Other professional or technical qualification, 3 NVQ or HND or HNC or equivalent, 4 Higher Grade, A levels, AS levels or equivalent, 5 Standard Grade, O levels, GCSEs or equivalent, 6 CSEs or equivalent, 7 - School leavers certificate, 8 - Other, 9 - No Qualification. All phenotypic data are standardized to mean zero and variance one.

### Simulation study

To demonstrate that our model is capable of accurately inferring phenotypic variations and correlations between multiple traits, we simulated epigenetic effects for two traits for the methylation data of chromosome 1 (*p* = 80, 545 probes), using three different scenarios for the epigenetic (co)variance matrix,

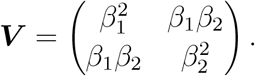

1. Scenario 1 represents a covariance matrix where there is no correlation between the two traits:

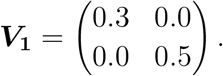
2. The second scenario introduces negative correlation between the two traits:

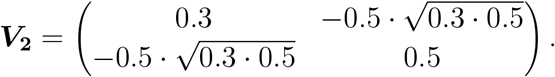
3. Scenario 3 assumes positive correlations between the two traits:

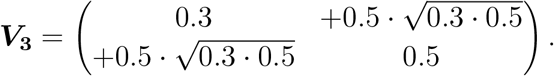

These matrices were scaled by the number of causal markers *p*_0_ = 1000 to sample the epigenetic effects from a multivariate normal distribution. When multiplying the simulated effects with their respective standardized columns of the **X** matrix, we obtained an epigenetic value, **g**, for each individual and trait. In each scenario, a vector of residuals was sampled from a normal distribution with variance (𝕀_*q*_ *− var*(***g***)) with the covariance elements set to 0, and added to **g** to obtain a matrix of phenotypes, **Y**. We repeated the data generation ten times for each of the three scenarios, where the causal effects were selected randomly. The datasets were then split into training data (n=17,264) and data (n=1000) for replication. The training data were anlaysed with MAJA, running the model for 2000 iterations. The posterior mean estimates of the effect sizes and their (co)variances were calculated using the last 1000 iterations. The posterior means of the effects covariances reproduce the true values very well for all three scenarios, as can be seen in Figure S1.

The estimated effect sizes 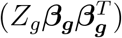, were then used to create predictors, 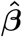, for each individual *i* in the test data to obtain the coefficient of determination

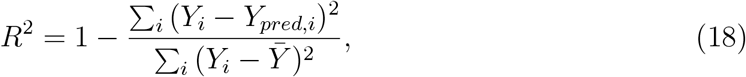

Where 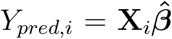 is the mean of generated phenotypes. Figure S3 shows that the estimated *R*^2^ agrees well with the expected *R*^2^ when the traits are correlated. The expected value is calculated according to Equation 34 in Ref.^7^ assuming *M*_*eff*_ = 30, 000 independent markers, a number estimated from the training data. The expected *R*^2^ is dependent on the assumed number of independent markers which is likely different for the case where the two traits are uncorrelated, which explains the large difference for estimated and expected *R*^2^ for *V*_1_.

We determined the true positive (TPR) and false discovery (FDR) rates of MAJA across simulation scenarios. True positives were identified as probes for which a causal effects was simulated, where the posterior inclusion probability was *≥* 0.95 and for which the posterior mean effect estimate was *≥ ±* 2 SD from zero. TPR was calculated as the number of true positives divided by the number of simulated causal variants. False positives were identified as probes that were not simulated to be causal variants, where the posterior inclusion probability was *≥* 0.95 and for which the posterior mean effect estimate was *≥ ±* 2 SD from zero. FDR was calculated as the number of false discoveries divided by the total number of discoveries. Figure S5 shows the TPR and FDR for the two simulated traits across scenarios.

Finally, to demonstrate that MAJA is also able to handle multiple groups and accurately estimate the covariances of each group, epigenetic effects and phenotypic information for two traits for the methylation data of chromosome 1 (*p* = 80, 545 probes) and chromosome 2 (*p* = 60, 707 probes) were generated. The effect sizes in the two chromosomes were generated according to three scenarios, where the second number in the subscript refers to the group (in this case chromosome):

1. 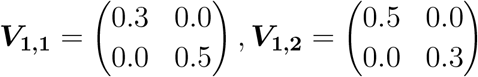
2. 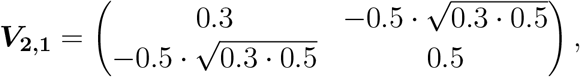
3. 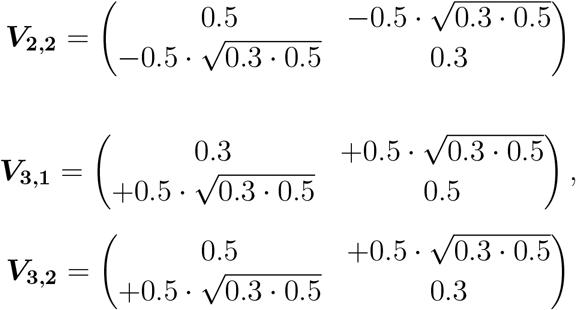

Each group was generated to have 500 causal markers. In each of the groups, the posterior means of the effects covariances reproduce the true values very well for all three scenarios, as displayed in Figure S2. Figure S4 shows that the estimated and expected *R*^2^ when the effects are estimated for two different groups with different covariances. The *R*^2^ values agree well when the traits are correlated.

### Prediction into the Lothian Birth Cohort

The Lothian Birth Cohort of 1936 (LBC1936) represents a longitudinal study of aging^12^. The 1091 cohort members were all born in 1936 and have been assessed for a wide variety of health and lifestyle outcomes. DNA has been collected at each clinical visit. In the present study, we considered DNA methylation data (Illumina 450k array) from whole blood, taken at mean age 70, for analysis. Details of the collection and processing of the data have been reported previously^18^. In brief, after quality control to remove poorly performing methylation sites, samples, and individuals with mismatching genotypes or predicted sex, a sample of 861 individuals was available for prediction analysis. The methylation and phenotypic data were processed in the same manner as GS. Additional phenotypes in LBC1936 were the Wechsler Test of Adult Reading; the National Adult Reading Test; and a general measure of cognitive function.

In LBC1936, BMI is calculated as weight in kilograms divided by height in meters. Weight and height were assessed at the wave 1 (baseline) clinic appointment. HDL cholesterol (mmol/L) and total cholesterol (mmol/L) are blood-based measurements from samples given in clinic at the baseline appointment. The cholesterol ratio is calculated as HDL cholesterol divided by total cholesterol. Scores for thirteen cognitive tests were available across five waves of data collection. Testing was performed triennially from age 70 to 82. Visuospatial ability was measured using the Block Design, Matrix Reasoning (WAIS-IIIUK) and Spatial Span (WMS-IIIUK) tests. Verbal ability was measured using the National Adult Reading Test, Wechsler Adult Reading Test and Verbal Fluency Test (using letters C, F and L). Memory was assessed via the Verbal Paired Associates, Logical Memory – a combination of immediate and delayed memory (WMS-IIIUK) and Digit Span Backwards (WAIS-IIIUK) tests. Processing speed was evaluated via the Digit Symbol Substitution Test, Symbol Search (WAIS-IIIUK), Choice Reaction Time and Inspection Time tests.

A latent measure of general cognitive function was obtained by using confirmatory factor analysis in a structural equation modelling (SEM) framework using the R package Lavaan (version 0.6-12)^19^. A first-order hierarchical cognitive model was specified.

Specifically, levels and change in general cognitive functioning were modelled with latent growth curve model (LGCM) using a Factor of Curves specification^20^. Intercepts and slopes of each cognitive test were used to indicate a latent intercept and slopes of general cognitive function and change. The growth curve slopes were weighted by mean lag time between each wave and baseline. Marker method was used to scale according to the first variable, and all models used full information maximum likelihood to include all data available. Negative residual variances were fixed to zero. Residual covariance between tests in the same cognitive domain were specified^21^.

Episcores were projected into LBC1936 wave 1 methylation data (n = 861). Linear regression was used to model each episcore (as a predictor) in relation to the outcome variables. Incremental *R*^2^ estimates are reported as the differences between models adjusting for age and sex compared to those that additionally include the episcore. For the variance explained in general cognitive function level, linear regression models were performed within Lavaan, with the G intercept from the latent growth curve models used as the outcome (see Ref.^13^). Model fit and test loadings can be found in Supplementary Table S5.

## Acknowledgements

We thank members of the Medical Geneomics group at ISTA for their comments, which improved this manuscript. This work was funded by an SNSF Eccellenza Grant to MRR (PCEGP3-181181), and by core funding from the Institute of Science and Technology Austria. We would like to acknowledge the participants and investigators of the Generation Scotland and Lothian Birth Cohort studies. Generation Scotland received core support from the Chief Scientist Office of the Scottish Government Health Directorates [CZD/16/6] and the Scottish Funding Council [HR03006]. Genotyping and methylation typing of the GS:SFHS samples was carried out by the Genetics Core Laboratory at the Wellcome Trust Clinical Research Facility, Edinburgh, Scotland and was funded by the Medical Research Council UK and the Wellcome Trust (Wellcome Trust Strategic Award “STratifying Resilience and Depression Longitudinally” (STRADL) Reference 104036/Z/14/Z). DNA methylation data for Generation Scotland was also funded by a 2018 NARSAD Young Investigator Grant from the Brain and Behavior Research Foundation (Ref: 27404; awardee: Dr David M Howard) and by a John, Margaret, Alfred and Stewart Sim Fellowship from the Royal College of Physicians of Edinburgh (Awardee: Dr Heather C Whalley). The LBC1936 is supported by the Biotechnology and Biological Sciences Research Council, and the Economic and Social Research Council [BB/W008793/1] (which supports SEH, and JC), Age UK (Disconnected Mind project), the Milton Damerel Trust, the Medical Research Council (G0701120, G1001245, MR/M013111/1, MR/R024065/1) and the University of Edinburgh. Methylation typing of LBC1936 was supported by the Centre for Cognitive Ageing and Cognitive Epidemiology (Pilot Fund award), Age UK, The Wellcome Trust Institutional Strategic Support Fund, The University of Edinburgh, and The University of Queensland. HS is supported by funding from the Wellcome Trust 4-year PhD in Translational Neuroscience [218493/Z/19/Z]. SRC is also supported by a Sir Henry Dale Fellowship jointly funded by Wellcome and the Royal Society [221890/Z/20/Z]. High-performance computing was supported by the Scientific Service Units (SSU) of IST Austria through resources provided by Scientific Computing (SciComp).

## Author contributions

IK and MRR conceived and designed the study. EB, MM, DLM and REM contributed to data preparation and design of the analyses. IK wrote the software and conducted the analyses, with assistance from HS for the prediction. REM, MRR, JC, SEH and SRC, provided study oversight. IK and MRR wrote the paper.

## Data availability

Access to the data is available with appropriate permission from the Generation Scotland Access Committee. Applications should be made to. (https://www.ed.ac.uk/lothian-birth-cohorts/data-access-collaboration) gives information on the Lothian Birth Cohort data, which are available on request from the Lothian Birth Cohort Study, University of Edinburgh. Data from both cohorts are not publicly available as they contain information that could compromise participant consent and confidentiality.

## Code availability

Source code is available at https://github.com/medical-genomics-group/MAJA.

## Supplementary information

**Table S1:** Estimated posterior (co)variances 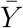, and 95% confidence intervals (CI) for body mass index (BMI), ratio of high density lipoprotein over total cholesterol (CHL), high blood pressure (BP), depression (DEP), osteoarthritis (OA) and asthma (AT).

| **variance** | **$\hat{V}$** | **95% CI** |
| --- | --- | --- |
| BMI | 0.8182 | 0.0270 |
| CHL | 0.7239 | 0.0339 |
| BP | 0.6877 | 0.0526 |
| DEP | 0.2426 | 0.0361 |
| OA | 0.5750 | 0.0777 |
| AT | 0.2367 | 0.0345 |
| BMI-CHL | -0.2939 | 0.0262 |
| BMI-BP | 0.1408 | 0.0274 |
| CHL-BP | -0.0306 | 0.0303 |
| BMI-DEP | 0.0692 | 0.0232 |
| CHL-DEP | 0.0102 | 0.0321 |
| BP-DEP | 0.0328 | 0.0392 |
| BMI-OA | 0.0721 | 0.0278 |
| CHL-OA | 0.0827 | 0.0397 |
| BP-OA | 0.2794 | 0.0488 |
| DEP-OA | 0.0017 | 0.0260 |
| BMI-AT | 0.0392 | 0.0245 |
| CHL-AT | -0.0096 | 0.0242 |
| BP-AT | -0.0679 | 0.0336 |
| DEP-AT | 0.0146 | 0.0106 |
| OA-AT | 0.0103 | 0.0384 |

**Table S2:** Estimated posterior (co)variances, 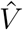, and 95% confidence intervals (CI) for years in education (EY), highest qualification in education (EQ), digit symbol (DS), logical memory (LM), verbal fluency (VT) and vocabulary (VO) tests.

| variance | $\hat{V}$ | 95% CI |
| --- | --- | --- |
| EY | 0.5791 | 0.0760 |
| EQ | 0.7778 | 0.0750 |
| DS | 0.6446 | 0.0475 |
| LM | 0.3232 | 0.0514 |
| VT | 0.4247 | 0.0647 |
| VO | 0.7311 | 0.1026 |
| EY-EQ | -0.5142 | 0.0927 |
| EY-DS | 0.3121 | 0.0436 |
| EY-LM | 0.2133 | 0.0480 |
| EY-VT | 0.1835 | 0.0542 |
| EY-VO | 0.3284 | 0.0986 |
| EQ-DS | -0.3799 | 0.0475 |
| EQ-LM | -0.2549 | 0.0613 |
| EQ-VT | -0.2371 | 0.0579 |
| EQ-VO | -0.4254 | 0.0882 |
| DS-LM | 0.2393 | 0.0368 |
| DS-VT | 0.1760 | 0.0425 |
| DS-VO | 0.1109 | 0.0527 |
| LM-VT | 0.0859 | 0.0483 |
| LM-VO | 0.1542 | 0.0623 |
| VT-VO | 0.3082 | 0.0579 |

**Table S3:**
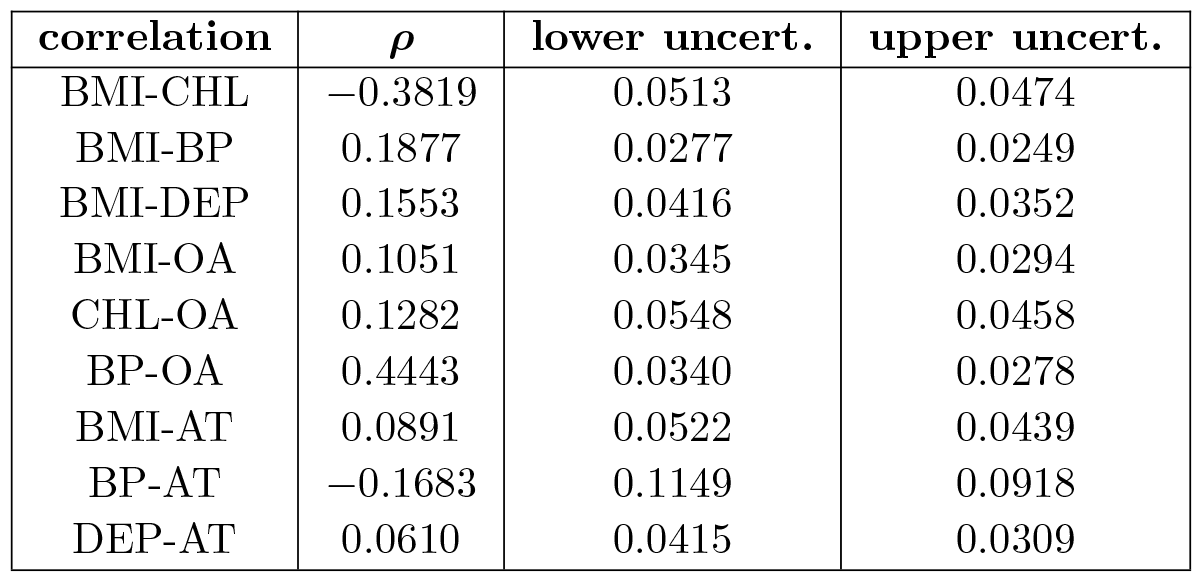
Correlations, *ρ*, calculated from the posterior (co)variances for body mass index (BMI), ratio of high density lipoprotein over total cholesterol (CHL), high blood pressure (BP), depression (DEP), osteoarthritis (OA) and asthma (AT), for the cases where the covariances were different from 0. The upper and lower uncertainties are calculated using the posterior means *±* 95% credible interval.

| **correlation** | **$\rho$** | **lower uncert.** | **upper uncert.** |
| --- | --- | --- | --- |
| BMI-CHL | -0.3819 | 0.0513 | 0.0474 |
| BMI-BP | 0.1877 | 0.0277 | 0.0249 |
| BMI-DEP | 0.1553 | 0.0416 | 0.0352 |
| BMI-OA | 0.1051 | 0.0345 | 0.0294 |
| CHL-OA | 0.1282 | 0.0548 | 0.0458 |
| BP-OA | 0.4443 | 0.0340 | 0.0278 |
| BMI-AT | 0.0891 | 0.0522 | 0.0439 |
| BP-AT | -0.1683 | 0.1149 | 0.0918 |
| DEP-AT | 0.0610 | 0.0415 | 0.0309 |

**Table S4:** Correlations, *ρ*, calculated from the posterior (co)variances for years in education (EY), highest qualification in education (EQ), digit symbol (DS), logical memory (LM), verbal fluency (VT) and vocabulary (VO) tests, for the cases where the covariances were different from 0. The upper and lower uncertainties are calculated using the posterior means *±* 95% credible interval.

| **correlation** | **$\rho$** | **lower uncert.** | **upper uncert.** |
| --- | --- | --- | --- |
| EY-EQ | -0.7661 | 0.2544 | 0.2022 |
| EY-DS | 0.5108 | 0.0209 | 0.0174 |
| EY-LM | 0.4930 | 0.0460 | 0.0344 |
| EY-VT | 0.3700 | 0.0661 | 0.0497 |
| EY-VO | 0.5046 | 0.0960 | 0.0731 |
| EQ-DS | -0.5366 | 0.1232 | 0.1038 |
| EQ-LM | -0.5083 | 0.2151 | 0.1658 |
| EQ-VT | -0.4125 | 0.1739 | 0.1351 |
| EQ-VO | -0.5640 | 0.2086 | 0.1642 |
| DS-LM | 0.5243 | 0.0215 | 0.0179 |
| DS-VT | 0.3364 | 0.0485 | 0.0391 |
| DS-VO | 0.1616 | 0.0666 | 0.0539 |
| LM-VO | 0.2318 | 0.1116 | 0.0815 |
| LM-VT | 0.3173 | 0.0949 | 0.0702 |
| VO-VT | 0.5531 | 0.0269 | 0.0200 |

**Table S5:** Loadings of cognitive tests on intercept. Latent measures of general cognitive function were generated using confirmatory factor analysis in a structural equation modelling (SEM) framework. Levels and change in general cognitive functioning were modelled with latent growth curve model (LGCM) using a Factor of Curves specification. The intercepts were used to indicate a latent intercept of general cognitive function and the test loadings are given. Model fit measures were calculated including confirmatory factor index (CFI = 0.958), Tucker-Lewis index (TLI = 0.957), root mean squared error approximation (RMSEA = 0.029) and the standardised root mean squared residual (SRMR = 0.061).

| Test | Intercept term |
| --- | --- |
| Block design | 0.702 |
| Matrix reasoning | 0.792 |
| Span total | 0.642 |
| NART | 0.685 |
| WTAR | 0.679 |
| Verbal fluency | 0.534 |
| Verbal paired associates | 0.543 |
| Logical memory | 0.615 |
| Digit backwards | 0.722 |
| Symbol search | 0.788 |
| Digit symbol | 0.668 |
| Inspection time | 0.491 |
| Four choice reaction time | 0.526 |

**Figure S1:**
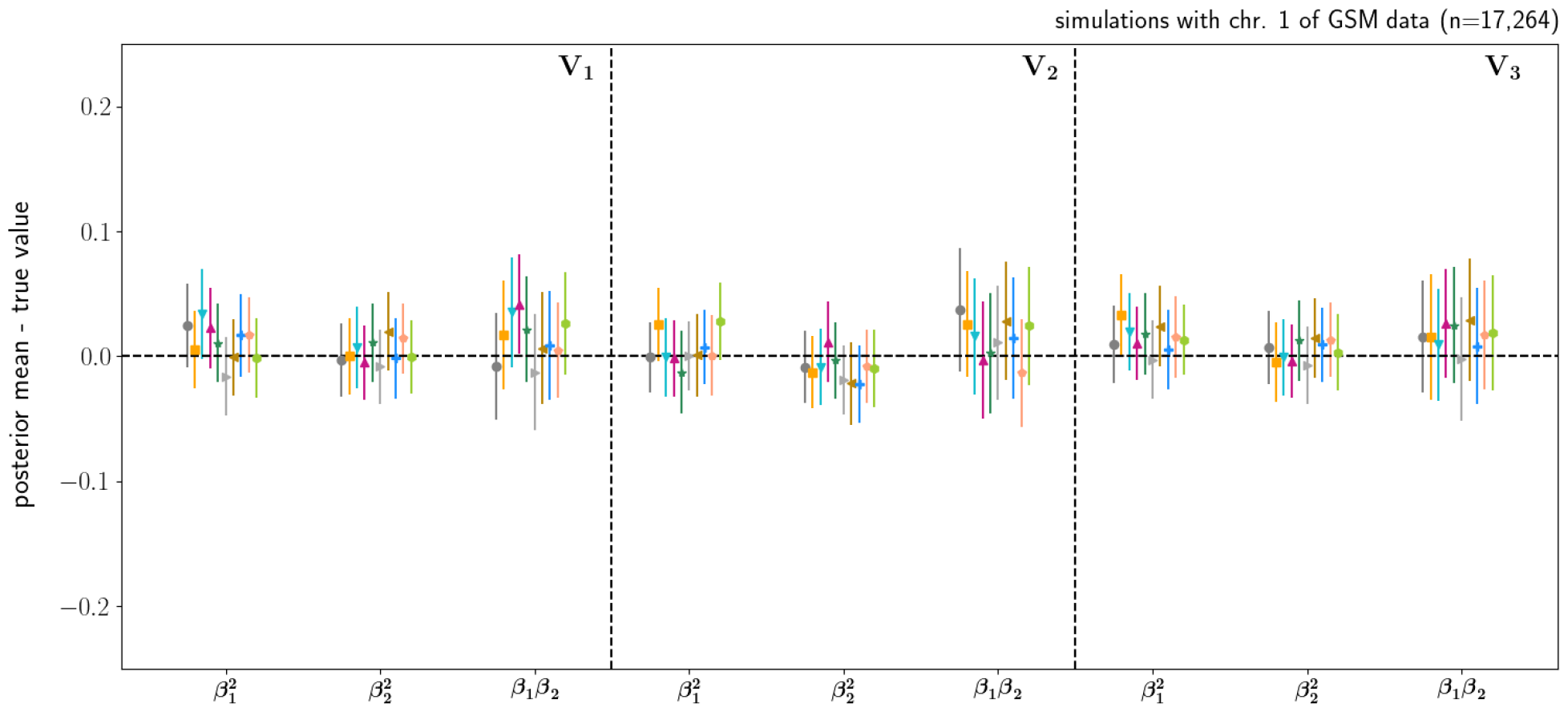
Simulation study results: Posterior mean estimates of the effects (co)variance components subtracted by their true value for each simulated dataset for three different covariance scenarios, denoted as *V*_1_ to *V*_3_. The error bars represent the 95% credible intervals.

**Figure S2:**
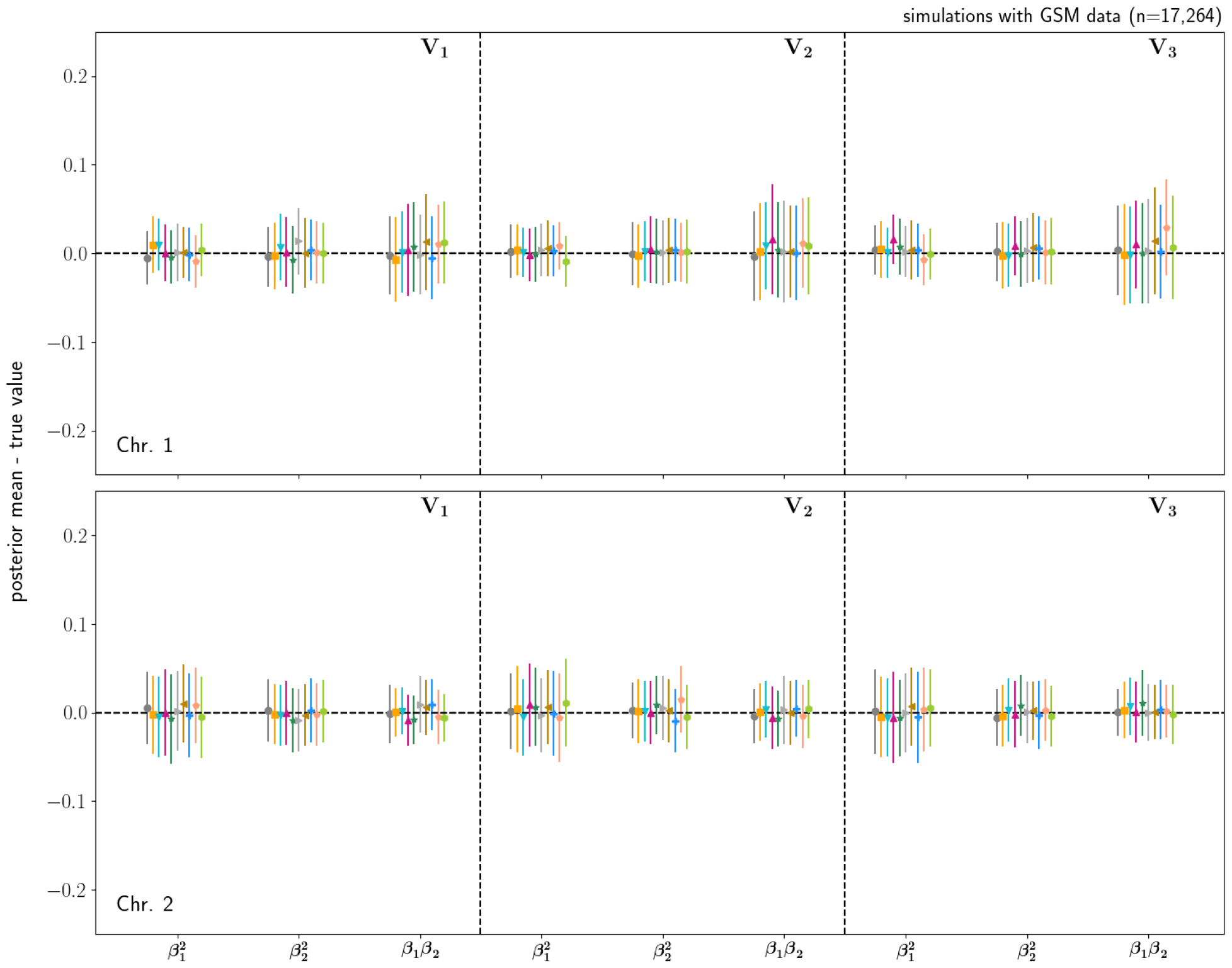
Simulation study results: Posterior mean estimates of the effects (co)variance components subtracted by their true value for each simulated dataset for three different covariance scenarios, denoted as *V*_1_ to *V*_3_, and two groups. The error bars represent the 95% credible intervals.

**Figure S3:**
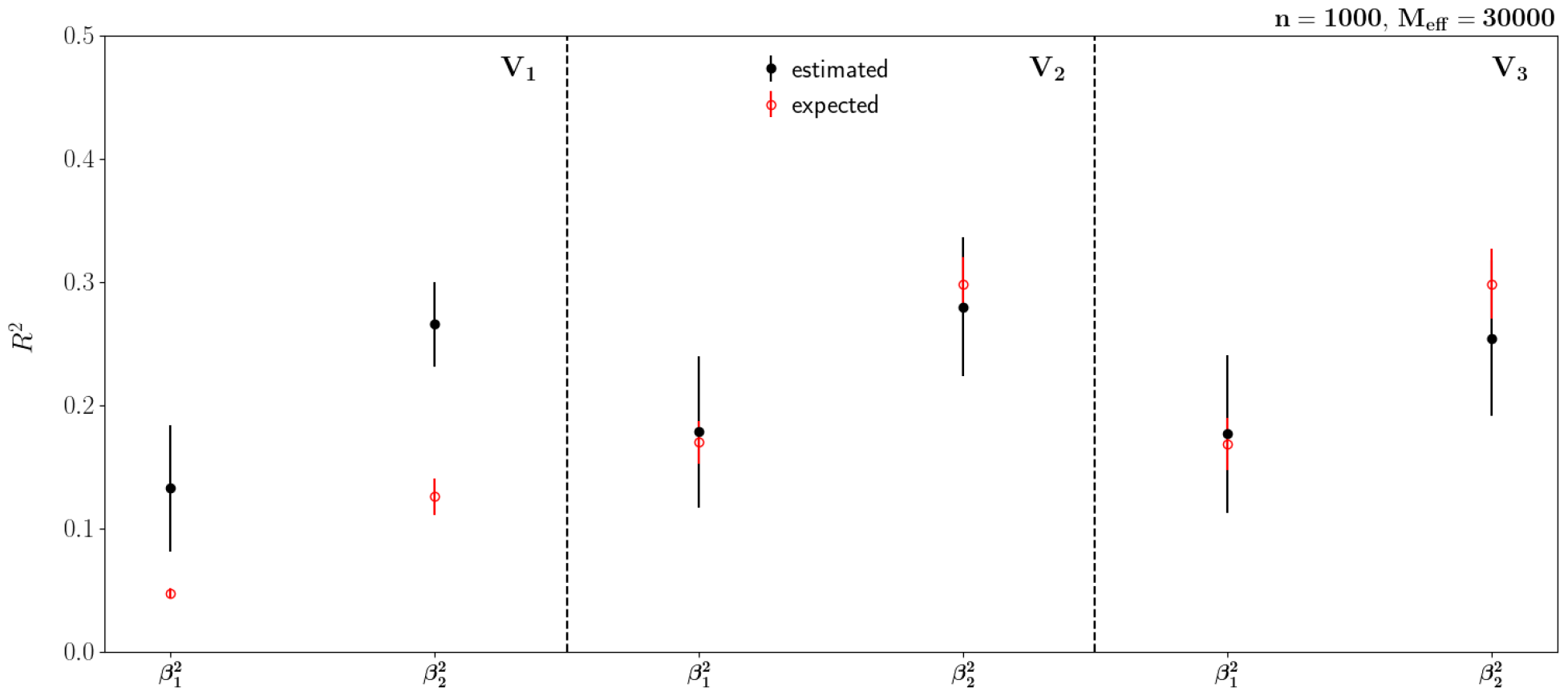
Mean expected and predicted coefficient of determination, *R*^2^, for 10 simulations and three covariance scenarios (*V*_1_ to *V*_3_) and a test dataset of 1000 individuals.

**Figure S4:**
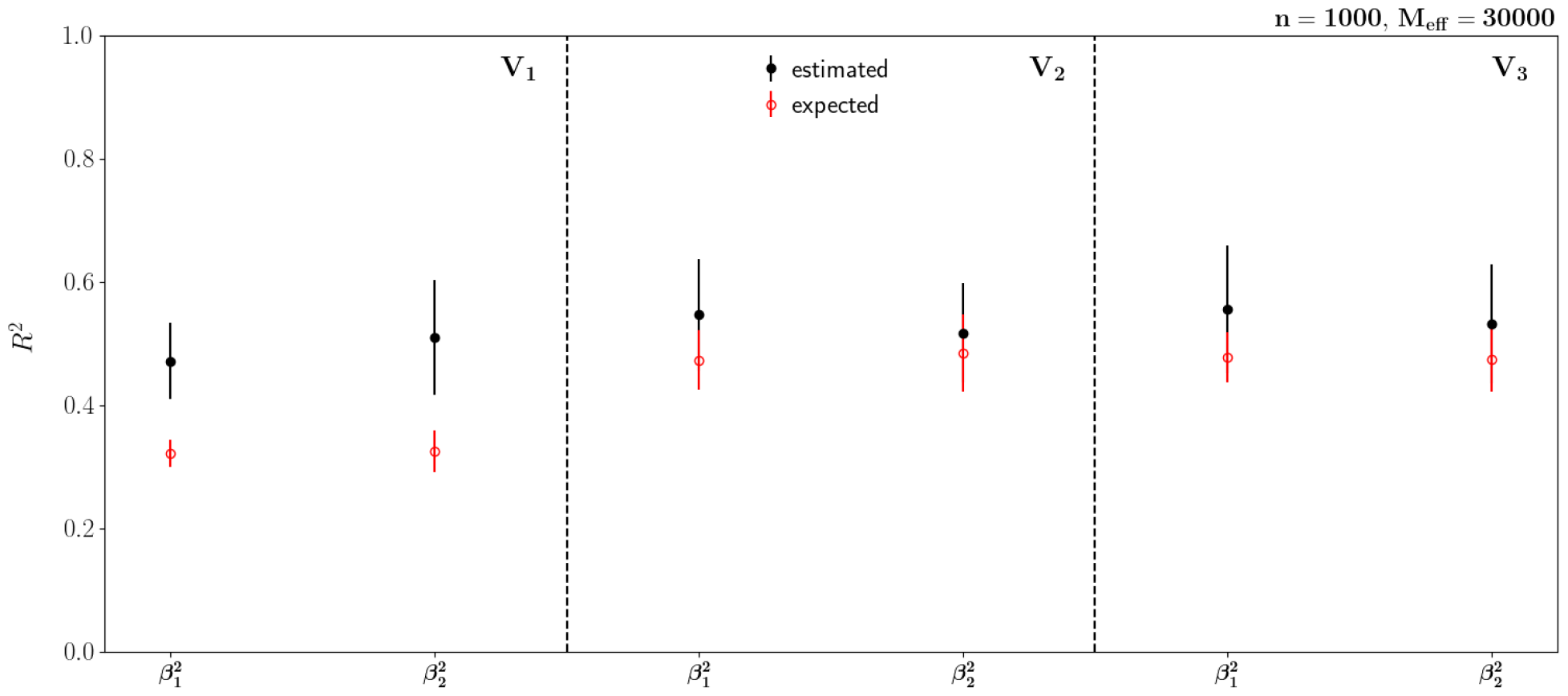
Mean expected and predicted coefficient of determination, *R*^2^, for 10 simulations and three covariance scenarios (*V*_1_ to *V*_3_) and a test dataset of 1000 individuals, where the effects were estimated for two different groups.

**Figure S5:**
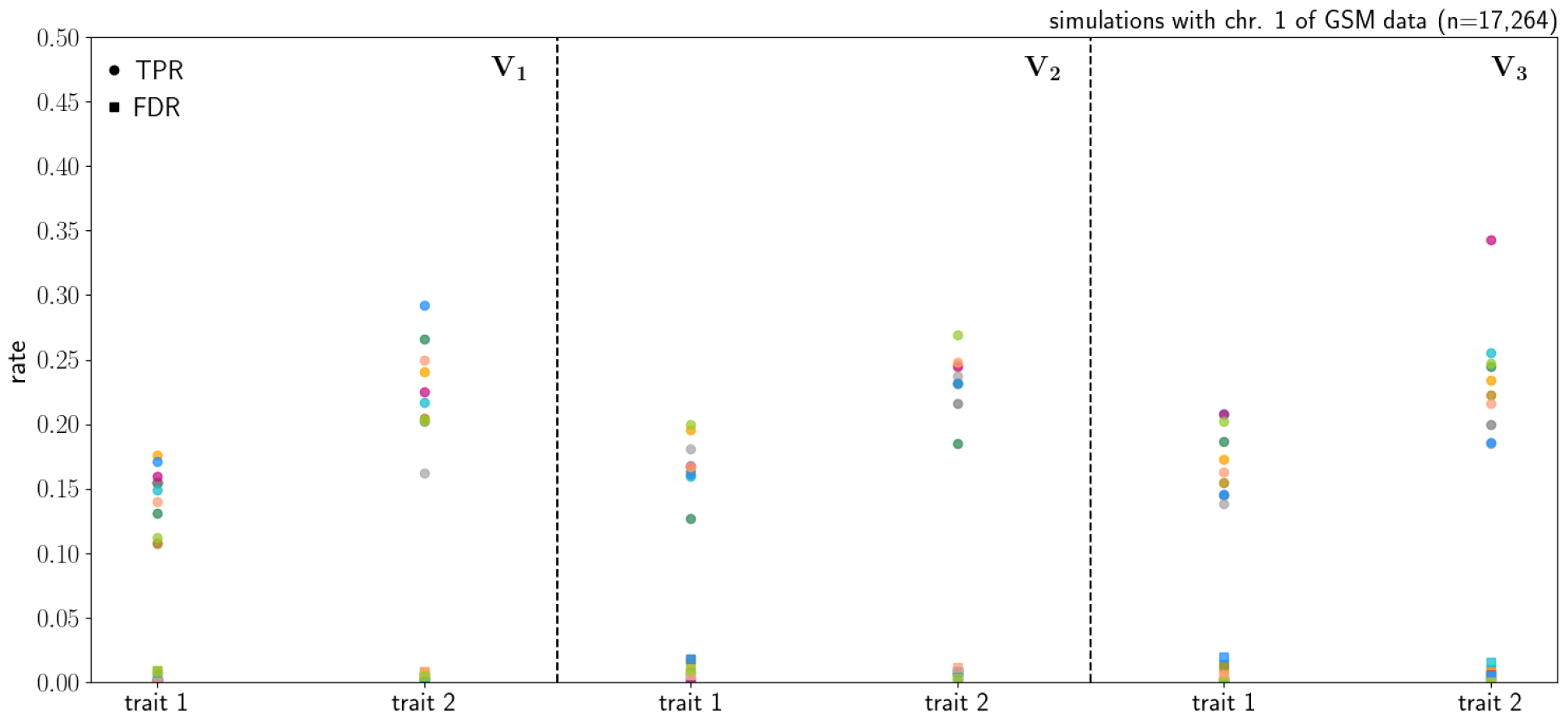
True positive rate (TPR) and false discovery rate (FDR) across 10 simulations (colours) for three different covariance scenarios, denoted as *V*_1_ to *V*_3_, between two traits. MAJA controls the FDR well below 2% for all scenarios, with the TPR (power) dependent upon the relationship among the traits.

**Figure S6:**
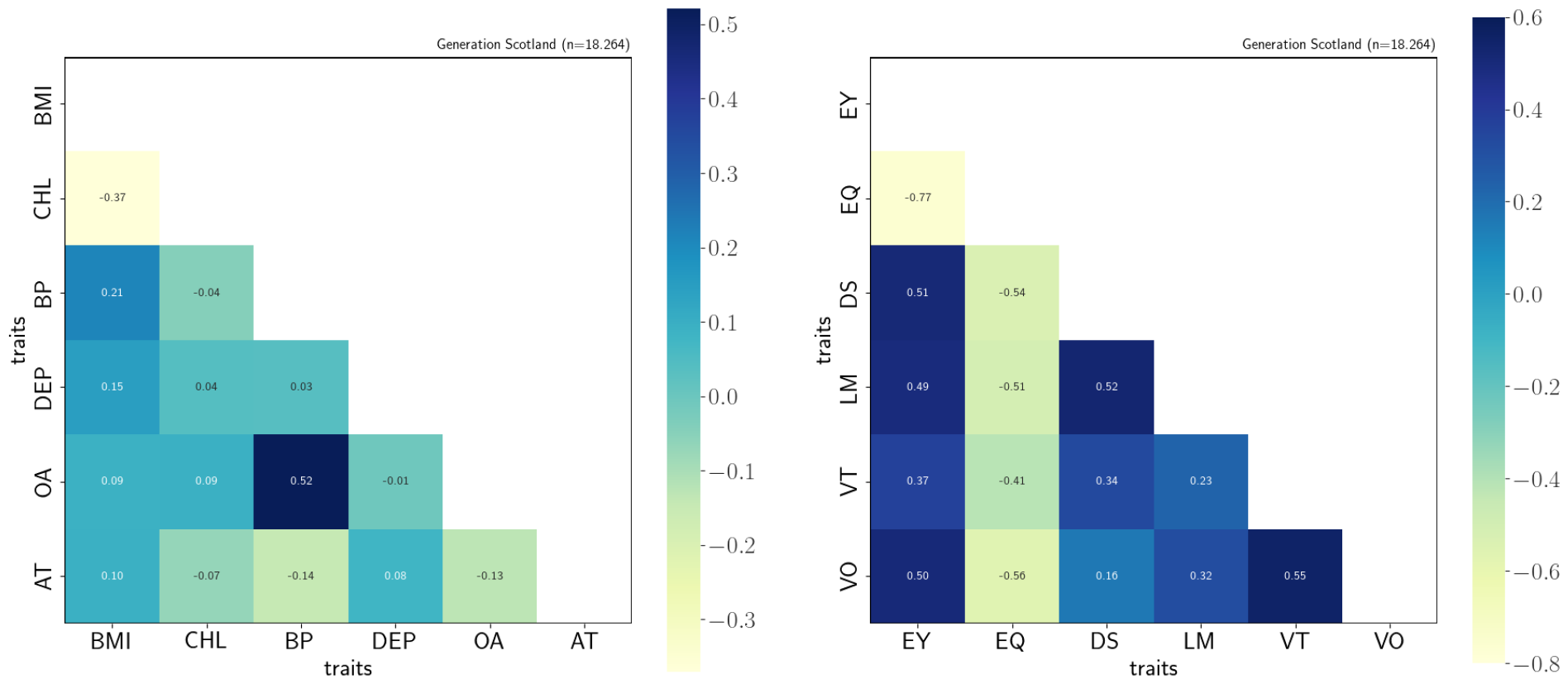
Correlations calculated from the posterior (co)variances for (left) body mass index (BMI), ratio of high density lipoprotein over total cholesterol (CHL), high blood pressure (BP), depression (DEP), osteoarthritis (OA) and asthma (AT) and (right) years in education (EY), highest qualification in education (EQ), digit symbol (DS), logical memory (LM), verbal fluency (VT) and vocabulary (VO) tests.

**Figure S7:**
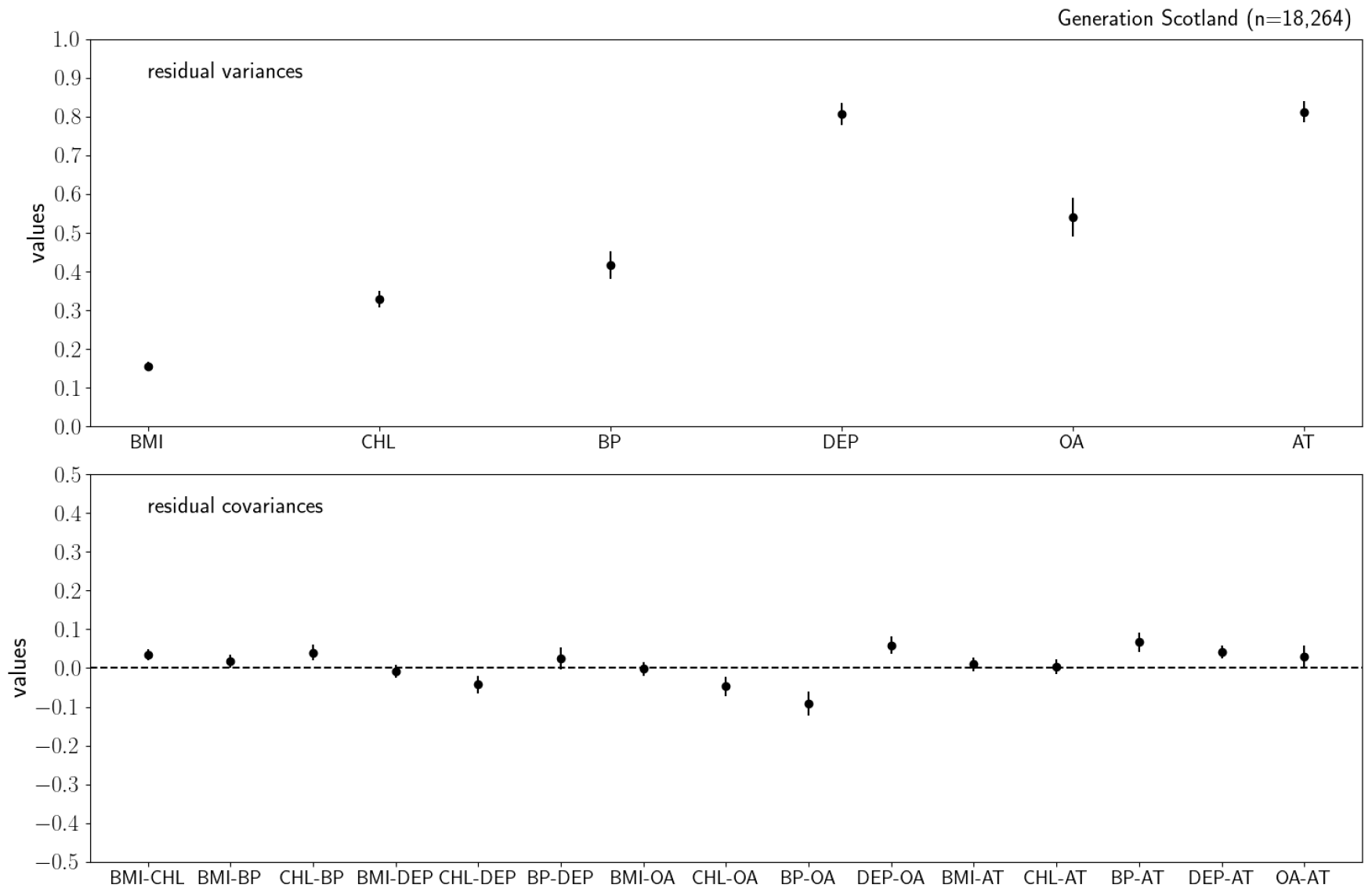
Estimated residual variances (top) and covariances (bottom) for body mass index (BMI), ratio of high density lipoprotein over total cholesterol (CHL), high blood pressure (BP), depression (DEP), osteoarthritis (OA) and asthma (AT) in the Generation Scotland methylation data using 18,624 individuals and 831,349 probes. The error bars represent the 95% credible interval.

**Figure S8:**
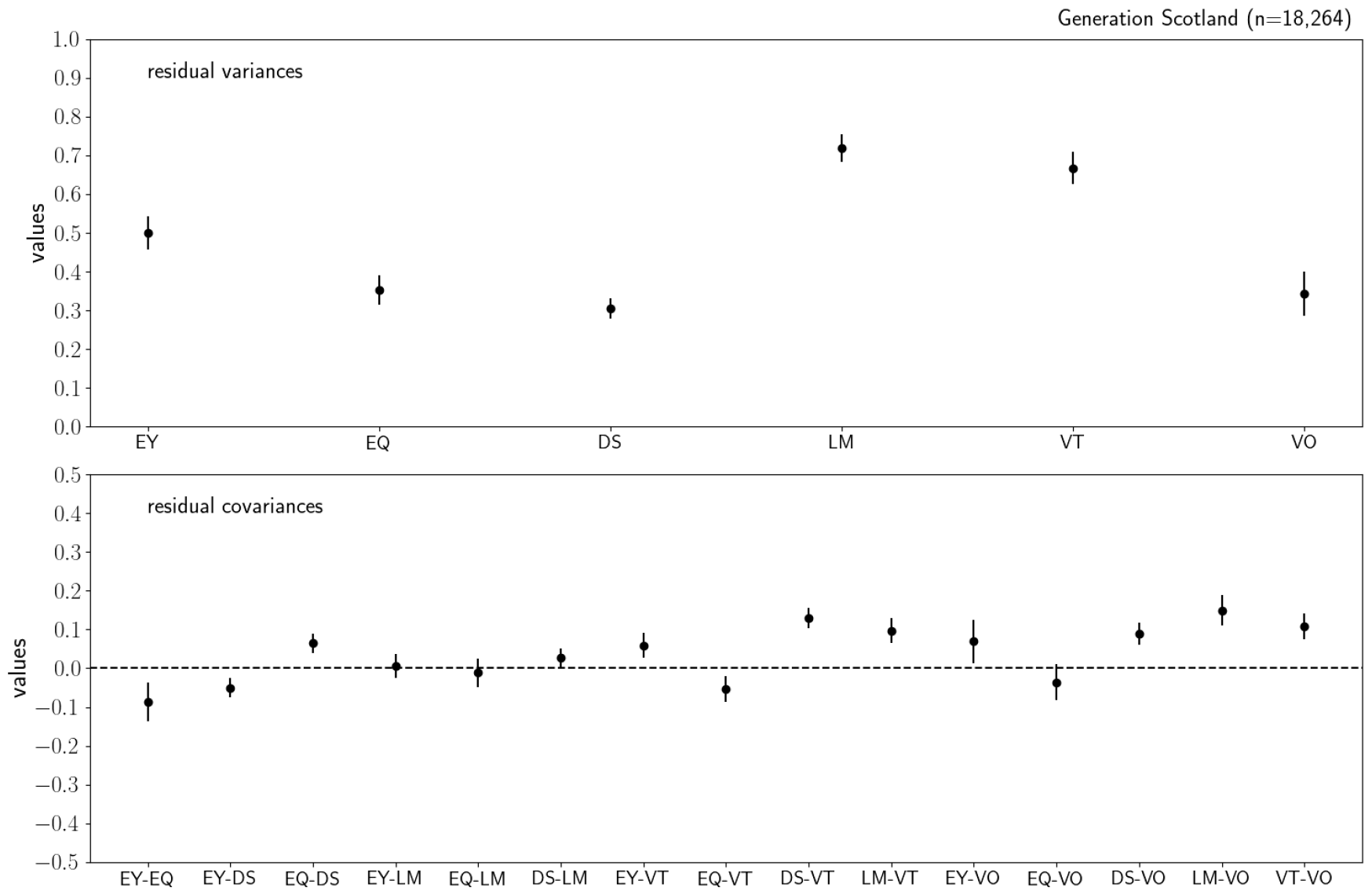
Estimated residual variances (top) and covariances (bottom) for years in education (EY), highest qualification in education (EQ), digit symbol (DS), logical memory (LM), verbal fluency (VT) and vocabulary (VO) tests in the Generation Scotland methylation data using 18,624 individuals and 831,349 probes. The error bars represent the 95% credible interval.

